# Neuronal endolysosomal acidification relies on interactions between transmembrane protein 184B (TMEM184B) and the vesicular proton pump

**DOI:** 10.1101/2025.02.01.635992

**Authors:** Elizabeth B. Wright, Erik G. Larsen, Marco Padilla-Rodriguez, Paul R. Langlais, Martha R.C. Bhattacharya

## Abstract

Disruption of endolysosomal acidification is a hallmark of several neurodevelopmental and neurodegenerative disorders. Impaired acidification causes accumulation of toxic protein aggregates and disrupts neuronal homeostasis, yet the molecular mechanisms regulating endolysosomal pH in neurons remain poorly understood. A critical regulator of lumenal acidification is the vacuolar ATPase (V-ATPase), a proton pump whose activity depends on dynamic assembly of its V0 and V1 subdomains. In this study, we identify transmembrane protein 184B (TMEM184B) as a novel regulator of endolysosomal acidification in neurons. TMEM184B is an evolutionarily conserved 7-pass transmembrane protein required for synaptic structure and function, and sequence variation in TMEM184B causes neurodevelopmental disorders, but the mechanism for this effect is unknown. We performed proteomic analysis of TMEM184B-interacting proteins and identified enrichment of components involved in endosomal trafficking and function, including the V-ATPase. TMEM184B localizes to early and late endosomes, further supporting a role in the endosomal system. Loss of TMEM184B results in significant reductions in endolysosomal acidification within cultured mouse cortical neurons. This alteration in pH is associated with impaired assembly of the V-ATPase V0 and V1 subcomplexes in the TMEM184B mutant mouse brain, suggesting a mechanism by which TMEM184B promotes flux through the endosomal pathway. Overall, these findings identify a new contributor in maintaining endosomal function and provide a mechanistic basis for disrupted neuronal function in human TMEM184B-associated nervous system disorders.

**Significance Statement:** Endolysosomal acidification is essential for neuronal protein homeostasis, yet its regulation in neurons remains poorly understood. Here, we identify TMEM184B as a key regulator of this process, establishing its first known cellular role. We show that TMEM184B interacts with vacuolar ATPase (V-ATPase) components and promotes the assembly of its V0 and V1 subdomains, facilitating lumenal acidification. Loss of TMEM184B disrupts endolysosomal pH in neurons, potentially impairing proteostasis. These findings reveal a critical function for TMEM184B in neuronal maintenance and provide mechanistic insight into its link to neurological disorders. This work advances our understanding of endolysosomal regulation and suggests TMEM184B regulation could improve outcomes in diseases involving lysosomal dysfunction.

## Introduction

Decreases in endolysosomal acidification contribute to cellular dysfunction in many neurological diseases. Cellular models of Lysosomal Storage Disorders (LSDs), Alzheimer’s Disease (AD), Parkinson’s Disease (PD), and Amyotrophic Lateral Sclerosis (ALS) all exhibit endolysosomal deacidification that causes the accumulation of toxic proteins (Arbo et al., 2020; Im et al., 2023; Hu et al., 2015; Lee et al., 2022; Lo & Zeng, 2023; Nixon & Rubinsztein, 2024; Root et al., 2021). In another disorder, Nieman Pick Type C Disease (NPCD), abnormally high pH in lysosomes causes failed trafficking of cholesterol cargos and neuron death (Lloyd-Evans et al., 2008; Vivas et al., 2019). Enlarged endosomal compartments or accumulation of autophagic vesicles in neurons occurs frequently in neurodegenerative diseases, implying that cargo delivery or degradation is impaired (Cataldo et al., 1996, 2000; Root et al., 2021). How endolysosomal deacidification initially occurs, and the mechanistic link between deacidification and disease pathogenesis, is not fully understood.

Endosomal trafficking relies on lumenal acidification mediated by the vesicular proton pump (V-ATPase). The V-ATPase consists of two subdomain complexes: the membrane-embedded V0 complex and the cytoplasmic V1 complex (Cotter et al., 2015; Forgac, 2007; Toei et al., 2010). When assembled, the V1 complex hydrolyzes ATP, driving the translocation of protons through a cytoplasmic hemichannel within the V0a subunit into the lumen. This proton transport gradually reduces the lumenal pH, facilitating compartment progression through the endolysosomal pathway.

The acidification of compartments is tightly regulated by the cell through dynamic assembly and disassembly of the V-ATPase in response to various factors, including amino acids, cholesterol, and intracellular signaling pathways (Colacurcio & Nixon, 2016; Hurtado-Lorenzo et al., 2006; Kane, n.d.; Ratto et al., 2022; Zoncu et al., 2011). Additionally, resident transmembrane proteins can modulate V-ATPase activity by sensing the availability of charged molecules to counterbalance proton accumulation in the lumen (Chadwick et al., 2021). For instance, the arginine transporter SLC38A9 is a well-characterized regulator of V-ATPase assembly, linking its activity to the activation of the mammalian target of rapamycin complex 1 (mTORC1) (Wyant et al., 2017). Under low-nutrient conditions, the V-ATPase complex is disassembled, leading to reduced mTOR activity and the induction of autophagy.

Transmembrane protein 184B (TMEM184B) is an evolutionarily conserved 7-pass transmembrane protein. TMEM184B is broadly expressed throughout the nervous system, with predominant expression in neurons (Bhattacharya et al., 2016; Larsen et al., 2022; Wright et al., 2023). Loss of TMEM184B leads to swollen presynaptic terminals, extra terminal branching, disrupted synaptic gene expression, and impaired synaptic transmission in multiple model systems, with concomitant disruptions in behavior (Bhattacharya et al., 2016; Cho et al., 2022). Electron microscopic analysis shows multilamellar inclusions within presynaptic terminals, suggesting a possible disruption in endolysosomal maturation, flux, or transport. Finally, TMEM184B human variants have been linked to a neurodevelopmental syndrome characterized by developmental delay, structural brain defects including corpus callosum hypoplasia and microcephaly, and seizures. In this study, TMEM184B patient-associated variants were linked to disruptions in cellular metabolism, as disruption of its function enhanced the nuclear localization of the stress-responsive transcription factor EB (TFEB) (Chapman et al., 2024). This finding suggests a possible role for TMEM184B in promoting cellular metabolism, which is interconnected with endolysosomal flux. Together, this evidence suggests that TMEM184B may contribute to neuronal structure and function by facilitating endolysosomal flux. How TMEM184B may accomplish this is unknown.

In this study, we investigated the functional role of TMEM184B in the endolysosomal system. We found that TMEM184B associates with proteins known to regulate intracellular transport and endolysosomal trafficking, including the V-ATPase. Analysis of endolysosomal acidification in the absence of TMEM184B revealed an elevated endolysosomal pH compared to wild-type controls, underscoring a functional role in endolysosomal function. We find reduced assembly of V-ATPase subcomplexes V0 and V1, which likely causes the pH disruptions in mutant neurons. These findings identify a mechanistic basis by which TMEM184B regulates neuronal endolysosomal acidification to ensure synaptic structure and function. Our data also suggest an explanation for how TMEM184B functional alteration disrupts neural development.

## Methods

### Experimental design and statistical analysis

For human cell TMEM184B localization, approximately 15-20 cells were imaged for each experimental group (TMEM184B-FLAG vs. GFP-FLAG) across 3 separate imaging sessions. For endolysosomal acidification assessment, a total of 13 mutant neurons and 10 wild-type primary cortical neurons were analyzed across 3 separate embryonic neuron dissections. Embryonic neurons were obtained from female mice at embryonic day 16. For V-ATPase assembly evaluation, 5 *Tmem184b*-mutant (3 male, 2 female) and 5 wild-type (3 male, 2 female) were euthanized for hippocampus dissection.

All statistical analyses were performed in GraphPad Prism, and p-values less than 0.05 were considered statistically significant. Statistical significance was defined as follows: p ≤ 0.05 (*), p ≤ 0.01 (**), p ≤ 0.001 (***). Data presented as the average ± SEM. For FIRE-pHly analysis, significance between wild-type and *Tmem184b-*mutant average puncta size was assessed using a Mann-Whitney test. All other analyses comparing *Tmem184b*-mutant and wild-type samples were evaluated by unpaired t-tests with Welch’s correction.

### Animals

The use of mice for this study has been approved by the University of Arizona IACUC (protocol 17-216). *TMEM184B*-mutant mice used in this study contain a gene-trap insertion originally created by the Texas A&M Institute for Genomic Medicine allele Tmem184bGt ^(IST10294F4)^ on the C57BL/6 background. Mice with this insertion exhibit < 5% of wild-type TMEM184B mRNA expression by both qPCR and RNAseq (Bhattacharya et al., 2016; Larsen et al., 2022). We therefore call this line a “mutant” rather than a “knockout” throughout the manuscript. C57BL/6 mice (littermates when possible) were used as wild-type controls throughout this study. All mouse husbandry was in accordance with guidelines of the Institutional Animal Care and Use Committee (protocol 17-216) at the University of Arizona.

### Reagents

Antibodies include: mouse monoclonal anti-V5-epitope antibody (Invitrogen, # R96025), myc-tag (9B11) mouse mAb (Cell Signalling Technology, # 2276S), rabbit polyclonal anti-ATP6V0A1 (Novus Biologicals, #NBP1-89342), rabbit polyclonal anti-ATP6V1A (Proteintech, # 17115-1-AP), rabbit polyclonal anti-ATP6V1H (abcam, # 187706), rat IgG Horseradish peroxidase-conjugated antibody (R&D systems, #HAF005), anti-rabbit IgG HRP-linked antibody (Cell Signalling Technology, #7074), monoclonal rat anti-LAMP1 (DSHB, # 1D4B), rabbit monoclonal anti-Cathepsin D (abcam, #75852), Cy3-goat anti-rabbit IgG (Jackson ImmunoResearch, # 111-165-144), Cy5-goat anti-rat IgG(Jackson ImmunoResearch, # 112-175-143), and purified anti-DYKDDDDK Tag antibody (BioLegend, #637301).

pcDNA4 CMV c-Myc-tag –GFP ORF –2xflag-tag and FCIV CMV BirA ORF-V5-tag IRES-Venus were ordered from Addgene. pcDNA3.1 TMEM184B vectors were ordered from GenScript. FLAG-hTMEM184B-GFP was generated by Twist Biosciences. FLAG-GFP was made in house. mCherry-Rab7a-7 was a gift from Michael Davidson (Addgene plasmid # 55127; http://n2t.net/addgene:55127; RRID: Addgene_55127). mCherry-Rab5a-7 was a gift from Michael Davidson (Addgene plasmid # 55126; http://n2t.net/addgene:55126; RRID: Addgene_55126). DsRed-rab11 WT was a gift from Richard Pagano (Addgene plasmid # 12679; http://n2t.net/addgene:12679; RRID: Addgene_12679). pLJM1-FIRE-pHLy was a gift from Aimee Kao (Addgene plasmid # 170775; http://n2t.net/addgene:170775; RRID: Addgene_170775).

### Cell culture and lysis

HEK293T cells were cultured in high glucose Dulbecco’s Modified Eagle Medium (DMEM; Gibco) containing 10% fetal bovine serum, 110mg/mL sodium pyruvate, and 10,000 U/mL Penicillin/Streptomycin. Cells were incubated at 5% CO_2_ at 37°C.

For immunoprecipitation coupled with tandem mass spectrometry (IP-MS) analysis and follow-up experiments, cells were removed from incubation and placed on ice for the entire lysis protocol. Media was aspirated from the plate and replaced twice with ice-cold PBS (Gibco) to gently wash cells. Cell scrapers (RPI) were used to pool cells for transfer to clean 1.5mL Eppendorf tubes. Cells were gently centrifuged at 800x g for 2 min. Cell pellets were resuspended with RIPA lysis buffer (150mM NaCl, 5mM EDTA, 50mM Tris, 1.0% Triton X-100, 0.5% sodium deoxycholate, 0.1% sodium dodecyl sulfate) containing EDTA-free protease inhibitor (Biomake). Cells were incubated at 4°C for 1 hour on a rotator followed by centrifugation at 21.1x g for 5 min. Supernatants were collected and passed through a 29 ga. Syringe (Exel International, 26028) seven times on ice prior to protein content quantification using a Pierce BCA protein assay kit (Thermofisher). Cells were stored at −80 °C for future IP-MS experiments.

### Primary cortical neuron culture

Cortices were collected from *Tmem184b-*mutant and wild-type mice at embryonic day 16. Neurons were dissociated with 0.25% Trypsin (Gibco) for 15 min at 37°C. Trypsin was replaced with 10% fetal bovine serum (FBS)-containing medium and incubated for 3 min at room temperature. Neurons were washed 3 times with ice-cold HBSS (Gibco) and triturated with fire polished glass Pasteur pipets (Fisher Scientific) of decreasing diameter 10 times each. Suspended neurons were filtered through 40µm cell strainers (Chemglass) and resuspended in Neurobasal Medium (Gibco) containing 0.5 mM L-glutamine, 10,000 U/mL Penicillin/Streptomycin (Gibco), and 1X B27 supplement. Approximately 20K neurons were plated in 8-well chambered #1.5 cover glass (Thermo Fisher) previously coated with 0.15ug/mL poly-D-lysine. Half of the media was replaced the next day prior to transfections. Neurons were incubated at 5% CO_2_ at 37°C.

### Transfections

Transfections in HEK293T cells were done using GeneJuice (EMD Millipore) and serum-free OptiMem GLUTAMAX (Gibco). For the entire IP-MS workflow, procedure was performed as previously described (Parker et al., 2019): expression vectors of TMEM184B variants were tagged with either the c-Myc epitope tag or the V5 epitope tag sequence. Control vectors included bifunctional ligase/repressor (BirA) tagged with the Myc tag or V5 tag. Each vector was then transfected into in duplicate.

To assess TMEM184B localization in the endosomal system, HEK293T cells were transfected with pHIV-FLAG-TMEM184B-myc or FLAG-GFP. To mark endosomes, cells were transfected with mCherry-Rab7 (late endosomes), mCherry-Rab5 (early endosomes), or dsRed-Rab11 (recycling endosomes)(Choudhury et al., 2002). Lysosome markers (CTSD and LAMP1) were evaluated using immunocytochemistry.

Transfections in primary cortical neurons were done using FuGENE HD transfection reagent (Promega) and OptiMem. After transfection with pFUGW-FIRE-pHly for pH analysis, neurons were incubated for 6 hr before media was changed. Imaging was performed 2 days after transfections.

### Immunocytochemistry

HEK293T cells were gently washed 3 times with 1X PBS-1% Tween 20 (PBST). Cells were fixed with 4% paraformaldehyde (PFA) in PBS for 15 min on ice. PFA was removed, cells were washed with 1X PBST for 5 min twice. Blocking solution (5% NGS in 1X PBST) was applied to cells and incubated for 30 min at room temperature on a rotator. Cells were gently washed 3 times with 1X PBST for 5 min to remove the remaining blocking solution. Primary antibodies against LAMP1 (1:500) and Cathepsin D (1:500) in 0.3% NGS and 1X PBST were applied to cells. Chambered slides were wrapped in parafilm and incubated overnight at 4°C. Primary antibodies were removed, and cells were gently washed 3 times with 1X PBST for 5 min. Secondary antibodies, goat, anti-rat Cy 5 (1:500) and anti-rabbit Cy3 (1:500) in 1X PBST, were added to cells. Chambered slides were covered in foil and incubated at room temperature for 1 hr. Cells were gently washed 3 times with 1X PBST for 5 min to remove any remaining secondary antibody and once with 1X PBS. Vectashield + DAPI (Fisher Scientific) was added to slides prior to applying #1.5 micro cover glass (Electron Microscopy Sciences)

### Hippocampus dissection

Mice were humanely euthanized with carbon dioxide, and hippocampi were removed within 30 min of euthanasia. Isolated hippocampi were immediately placed into RIPA lysis buffer with protease inhibitor. Hippocampi were crushed using clean pestles and sonicated before centrifugation at 21k x g for 5 min. Protein concentrations from the final supernatant were quantified via BCA. Hippocampi were collected from 5 *Tmem184b*-mutant and 5 wild-type controls. All mice were approximately 6 months old.

### Immunoprecipitation

Constructs containing mouse TMEM184B (NP_766196.1, 407 amino acids) or human TMEM184B (NP_036396.2, 407 amino acids) were used as appropriate. Mouse and human TMEM184B sequences are 96% identical (391/407) and 97% similar (395/407). 500 ug of Protein A/G Magnetic Beads (MedChemExpress, HY-K0202) were loaded into 1.5 mL Eppendorf tubes and washed 3 times with PBS prior to use. Beads were conjugated with primary antibody with incubation for 2 hr at room temperature on a rotator. Bead-antibody conjugates were washed 4 times with 1X PBS before addition of lysates and incubated overnight at 4°C on a rotator. Lysate supernatants were removed, and antibody-antigen were washed 4 times with PBS and transferred to new 1.5mL Eppendorf tubes to avoid non-specific binding to any remaining lysates. Proteins were eluded twice with 1X Laemmli sample buffer (Cold Spring Harbor) for a final volume of 40ul. Eluents were incubated for 5 min (95°C or 37 °C for TMEM184B samples) prior to loading into gel.

For mass spectrometry analyses, 5µg of anti-V5-epitope antibody (Invitrogen) was used for bead conjugation. 1.5 mg whole cell lysates were added to conjugated beads prior to incubation overnight. For V-ATPase assembly analysis, 0.16 µg anti-ATP6V0A1 primary antibody was used to conjugate beads. 100 µg of hippocampus protein was added to conjugated beads.

### Gel electrophoresis, Coomassie, and Immunoblotting

For V-ATPase assembly analysis, whole hippocampus lysates were mixed with 1X Laemmli Sample Buffer and incubated for 5 min at 95°C. Whole lysate controls and IP eluents were loaded into 4-20% Mini-PROTEAN TGX protein gels (Bio-Rad). Proteins were transferred onto 0.45m PVDF transfer membranes (Thermo Scientific) and blocked with 5% casein in 1X TBST (Sigma-Aldrich). Membranes were incubated overnight at 4°C in primary antibody against ATP6V1H or ATP6V1A in 0.3% casein and 1X TBST. Primary antibodies were removed, and membranes washed before incubation with horse-radish peroxidase-conjugated antibody. Blots were developed using ECL blotting substrate (Bio-Rad) and quantified using Image Lab software (Bio-Rad). For IP samples, V1A and V1H quantification was normalized to the total amount of V0A1 in the IP lane. Whole cell lysate controls were normalized to total amount of protein loaded into the sample lane.

For IP-MS, eluents were run fresh on the day of MS analysis. IP-MS gels were stained with mass-spectrometry grade Coomassie Blue, imaged for annotation purposes, and processed and analyzed by the University of Arizona College of Medicine’s Quantitative Proteomics Laboratory.

### Mass Spectrometry Data Acquisition and Analysis

To identify TMEM184B true protein-protein interaction candidates, a multiple epitope tags co-IP approach was used: two separate plasmids were created containing the same TMEM184B protein coding sequence but with either a c-Myc tag sequence or a V5 tag sequence at the end of TMEM184B’s C-terminus. In parallel, HEK293T 100mm plates were transfected with one vector in duplicate– thus, 8 plates in total were transfected: duplicates of TMEM184B-c-Myc, TMEM184B-V5, GFP-c-Myc, and BirA-V5. Mass spectrometry data was acquired as previously described (Kruse et al., 2017; Parker et al., 2019). Once acquired, Scaffold software-derived (version 5, Proteome Software Inc., Portland, OR) total spectrum counts (TSCs) were compiled into a CSV and uploaded into the RStudio GUI (version 4.1.0) using the 64bit R software (version 4.0.4). Upon uploading Scaffold-mediated TSC CSV files into R, the data was wrangled to enable mathematical operations on TSCs. The only departure from the methods above was a lack of deployment of SAINT scoring and analysis of SAINT-scored hits (Choi et al., 2010).

To analyze the resulting MS data, we filtered to remove low abundance proteins (< 5 TSCs in both TMEM184B IP duplicates and < 10 TSCs in every TMEM184B sample) (Teo et al., 2013)). Following filtering, subsequent processing was performed on each tag dataset separately. Values of 0.1 were added to proteins with 0 TSCs to enable fold change calculation(Choi et al., 2010; Sardiu et al., 2008; Sowa et al., 2009). TSCs were then normalized by molecular weight (MW) and averaged between duplicates. These averages were used to calculate fold changes (FCs), one for each tag (e.g., avg TMEM184B-c-Myc MW-normalized TSC / avg GFP-c-Myc MW-normalized TSC), and a combined FC (e.g., avg of all TMEM184B MW-normalized TSC / avg of all control bait MW-normalized TSC). Candidates with ≥ 2 Avg FC in both tag system and a ≥ 2 combined FC were retained, and common contaminants were filtered out as previously described (Mellacheruvu et al., 2013). R scripts, markdowns, and notebooks were developed to adapt multiple algorithms, to analyze subsequent candidates via gene ontology, and to visualize results.

### Fluorescent microscopy

All imaging was taken on the Nikon Spinning-Disk SoRa super-resolution microscope in the University of Arizona Cancer Center (UACC) Microscopy Shared Resource. Images were acquired on a Nikon CSU-W1 SoRa Spinning-Disk Confocal microscope equipped with a Photometrics Kinetix sCMOS camera. Single slice images were acquired using a Nikon 60x Plan Apo 1.40NA objective lens.

Prior to imaging live primary neurons, neurobasal media was removed from 8-well cover glass chambers and replaced with pre-warmed Live Cell Imaging Solution (Thermo Fisher). Neurons were allowed to acclimate to the microscope live cell imaging chamber (5% CO_2_ and 37°C) for 30 minutes prior to imaging. The 488 and 561 nm lasers were used to excite mTFP1 and mCherry, respectively. Single slices were obtained and exported as ND2 files for later analysis in Nikon NIS Elements AR. Wild-type (10) and Tmem184b-mutant (13) neurons were imaged across 3 separate groups of 1-2 pooled embryos per genotype.

For HEK293T localization experiments, the 405, 488, and 640 nm lasers were used to excite DAPI, GFP, and Cy5, respectively. The 561nm laser was used to excite mCherry, dsRed, or Cy3 according to the specific compartment. Single slice images were acquired for analysis in ImageJ (FIJI). For each compartment assessed, 15-20 images were taken for TMEM184B and GFP controls each.

### Colocalization analysis

Raw CZI files were imported into ImageJ (FIJI) for processing, including background subtraction, prior to analysis. To assess the co-localization of TMEM184B with other cellular compartments, images from the red and green channels were binarized and segmented using the watershed function. Overlapping puncta between the binarized images were identified and analyzed using the “AND” function in the Image Calculator. Both the merged puncta and individual puncta from each channel were quantified using the “Analyze Particles” function to determine puncta area (µm²). The degree of TMEM184B localization to each compartment was calculated by dividing the area of merged puncta by the total area of TMEM184B+ puncta. For lysosome localization, images labeled with CTSD (Cathepsin D) and LAMP1 (Lysosomal-associated membrane protein 1) were first processed using the “AND” function to identify puncta associated with lysosomes. This resulting image was then intersected with TMEM184B+ puncta to assess co-localization. Additionally, separate calculations for LAMP1 and TMEM184B were performed to examine TMEM184B localization to late endosomes or deacidified lysosomes, distinguishing these from conventional lysosomal compartments.

### FIRE-pHly puncta analysis

Nikon NIS Elements AR 5.42.03 software with the General Analysis 3 (GA3) module was used for image processing and analysis. To quantify red channel detections within each cell, a new image channel was created by applying a gaussian filter (sigma = 8) and rolling ball background subtraction (radius = 18µm) using the green channel. A signal threshold was applied to this “Cell Body” channel to properly segment the cell body and only quantify the red channel detections within each cell. The following image preprocessing tools were used for both the green and red channels: low pass filter = 4px and rolling ball background subtraction radius = 1.2µm. The Bright Spots Detection tool (diameter = 1.5µm) was used to detect the red channel within the Cell Body and the intensity threshold was adjusted per image to achieve accurate segmentation. The following data was quantified for each cell: cell body area (µm^2^), total channel detections, and mean intensity of green and red fluorescence. Weights were determined as follows: dividing each cell’s number of puncta by all the puncta in the study and then multiplying each cell’s fraction by 10 to re-scale the value of the weight near a whole mouse. The green-to-red (G/R) fluorescence ratio for each identified puncta was calculated by dividing the mean green fluorescence intensity by the mean red fluorescent intensity.

## Results

### TMEM184B interacts with multiple proteins involved in endosomal trafficking and autophagy

To clarify the role of TMEM184B in cellular processes that could contribute to neuronal maintenance (Bhattacharya et al., 2016; Cho et al., 2022), we first sought to identify its protein interactors within human cells. Because currently available antibodies show nonspecific binding in mutant tissue (M.R.C.B., personal communication) we could not cleanly immunoprecipitate endogenous TMEM184B from human or mouse tissue. We therefore performed immunoprecipitation (IP) of expressed, tagged TMEM184B coupled with tandem mass spectrometry (IP-MS/MS) analysis in HEK293T cells. We performed IP with two independent epitope tags (V5 and Myc) in duplicate for each tag to improve confidence in hits (Fig. 1A-B). After filtering out common contaminants (Mellacheruvu et al., 2013) and intersecting potential candidate interactors from both datasets, we identified 136 unique proteins as candidate interactors of TMEM184B (Fig. 1C and Extended Data Table 1-1).

**Fig 1:**
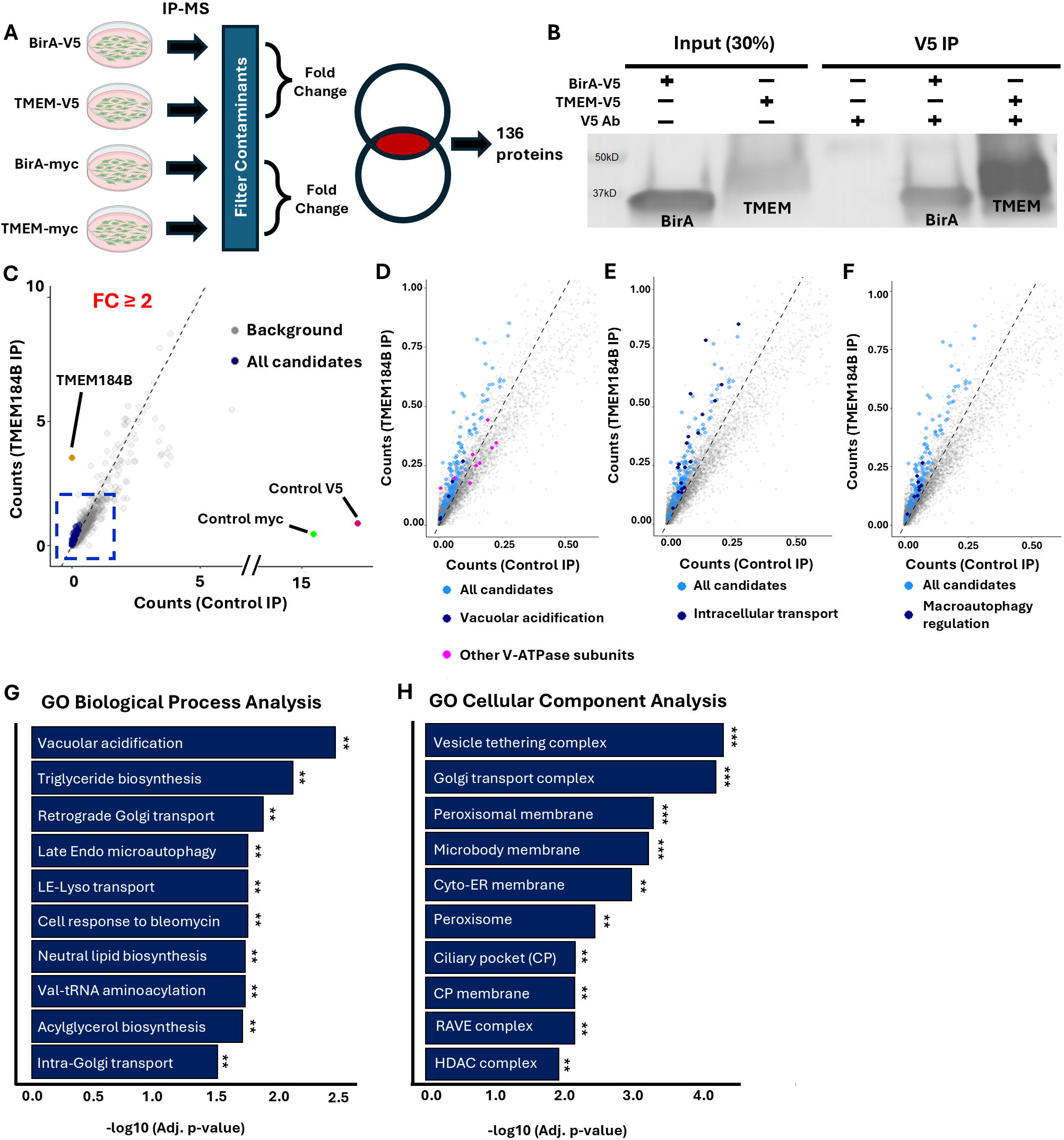
TMEM184B interacts with multiple proteins involved in endosomal trafficking and autophagy. **A**, Schematic depicting overall workflow of IP-MS analysis. **B,** Example western blot showing a successful pulldown of TMEM184B targeting its V5 tag. **C,** Average spectral counts of TMEM184B versus control (myc or V5) immunoprecipitation from HEK293T cells. Data is from 4 independent IP-MS experiments using two different affinity tags with 2 replicates each. Blue represents potential candidates significantly enriched as interactors with TMEM184B. All other proteins are denoted as “Background” in gray, and “TMEM184B” is marked in gold. Dashed box represents area in which all 136 enriched proteins are located. **D-F,** Cropped scatterplots (from blue dashed box in C) highlighting enriched proteins in functional categories. Light blue represents all protein candidates and dark blue represents proteins within designated functional category. Pink in D represents other vacuolar acidification GO term members. **G-H,** Top 10 significant results from gene ontology biological process (G) and cellular component (H) analyses, sorted by fold enrichment and adjusted p-value of 136 significant IP-MS hits.

Interestingly, TMEM184B demonstrated significant interactions with two isoforms of the V-ATPase subunit V0a (a1 and a2) (FC = 3.38 and 3.18, respectively). To confirm a potential interaction between FLAG-TMEM184B-GFP and ATP6V0A1 (the V0a1 subunit), we performed co-IP assays (Extended Data Figure 1-2). We first used the FLAG-tagged proteins as bait and probed for V0a1. Results confirmed an interaction between V0a1 and TMEM184B We performed an inverse co-IP using V0a1 as bait and probed for FLAG-tagged proteins to increase confidence in this interaction and found similar results. Regulators of V-ATPase assembly were also found among the interacting protein candidates, including the Drosophila melanogaster X chromosomal gene-like proteins (DMXL1 and DMXL2), also known as Rabconnectin-3 (FC = 4.05 and 4.44, respectively) (Eaton et al., 2024). Together, these data suggest that TMEM184B and the V-ATPase are associated in human cells.

Among hits in the mass spectrometry analysis, many proteins were involved in cellular trafficking or autophagy, suggesting a possible influence of TMEM184B in these dynamic processes (Extended Data Table 1-1). Notably, proteins involved in regulating endosomal trafficking to the plasma membrane include ADP-Ribosylation Factor Guanine Nucleotide Exchange Factor 2 (ARFGEF2, FC = 4.72), AAA-ATPase VPS4B (FC = 3.25), Coiled-Coil Domain Containing 22 (CCDC22, FC = 3.43), and the EH Domain Containing protein family 3 (EHD3, FC = 3.35) (Bar et al., 2013; Singla et al., 2019; Tseng et al., 2021; Zhu et al., 2022). Collectively, these proteins contribute to endosomal trafficking, each fulfilling a distinct function in vesicle trafficking, protein recycling, and the maintenance of cellular homeostasis. Autophagy-related proteins Tuberous Sclerosis Complex (TSC2, FC = 2.55) and Activating molecule in BECN1-regulated autophagy protein 1 (AMBRA, FC = 2.433) also were present among TMEM184B interactors. These proteins regulate autophagy through distinct mechanisms: TSC2 modulates mTORC1 and AMPK signaling pathways, and AMBRA1 directly participates in autophagosome formation (Albishi, 2023; Di Bartolomeo et al., 2010; Li et al., 2022; Nardo et al., 2014; Ng et al., 2011).

To identify enriched functional categories of TMEM184B-interacting protein candidates, we performed Gene Ontology (GO) analyses for Biological Processes (BP) and Cellular Components (CC). Multiple categories indicated involvement in vacuolar acidification, encompassing the V-ATPase and known regulators (fold enrichment = 20.71) and intracellular transport (fold enrichment = 2.48) (Fig. 1F). For CC enrichment, the Regulator of the H+-ATPase of vacuolar and endosomal membranes (RAVE) complex (Jaskolka, Tarsio, et al., 2021; Jaskolka, Winkley, et al., 2021; Smardon et al., 2002) showed the highest fold enrichment (99.42, FDR = 0.008) (Fig. 1G). Other significantly enriched GO categories included transport-related mechanisms, such as vesicle transport, membrane fusion, and Golgi trafficking (Extended Data Table 1-3). These results suggest that TMEM184B localizes to multiple organellar membranes.

### TMEM184B localizes to early and late endosomes in human cells

TMEM184B colocalizes with recycling endosomes in mouse sensory neurons and in fly motor neurons (Bhattacharya et al., 2016; Cho et al., 2022), but its distribution in human cells has not been established. Because many of the TMEM184B interactors we identified reside on endosomal membranes or participate in endosomal trafficking (Fig. 1D-E), we hypothesized that TMEM184B may at least partially localize to these compartments. We first quantified the fraction of TMEM184B^+^ puncta that overlapped with the endosomal markers Rab5 (early endosomes), Rab7 (late endosomes), and Rab11 (recycling endosomes) relative to the total population of TMEM184B^+^ puncta in each cell. Of all TMEM184B puncta identified, 45% showed early endosome colocalization, 34% showed late endosome colocalization, and 13% showed recycling endosome colocalization. (Fig. 2A-C, E). We further examined the prevalence of TMEM184B on each type of endosome. TMEM184B is present on ∼73% of early endosomes, ∼59% of late endosomes, and ∼45% of recycling endosomes. There is a small amount of TMEM184B at the plasma membrane, likely resulting from recycling endosome fusion. Together, these data indicate that TMEM184B is broadly distributed across endosomal populations, consistent with IP-MS results.

**Fig 2:**
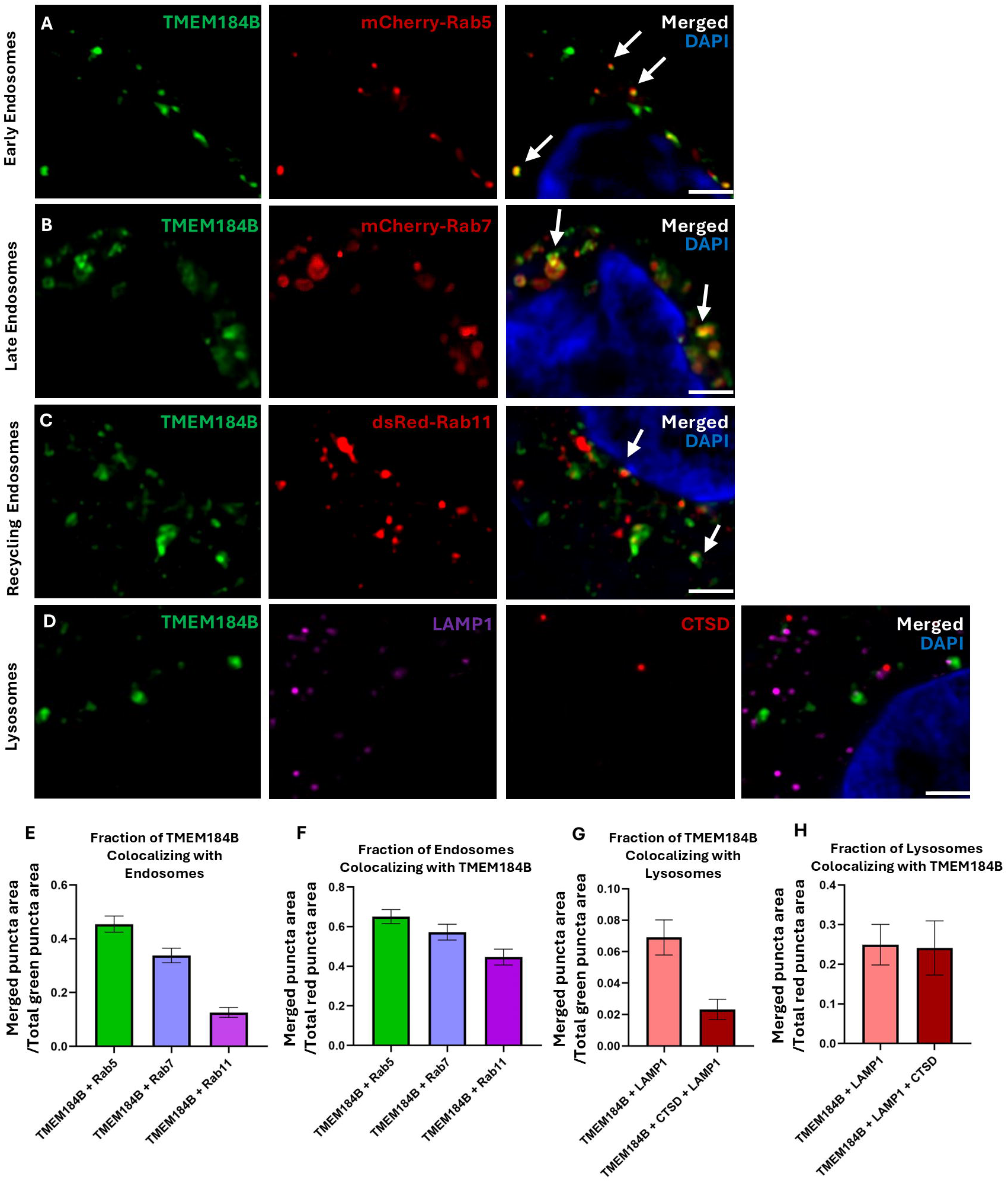
TMEM184B localizes to early and late endosomes in human cells. All images show HEK293T cells expressing FLAG-TMEM184B-GFP. DAPI (blue) marks the nucleus. Images represent single slices of human cells. **A-D,** images depicting localization of FLAG-TMEM184B-GFP to early endosomes (mCherry-Rab5, A), late endosomes (mCherry-Rab7, B), recycling endosomes (dsRed-Rab11, C), and lysosomes (LAMP1+CTSD, D). White arrows highlight overlapping puncta. Scale bars = 2 μm. n = 45-60 cells per marker, imaged over 3 sessions each. **E,** Fraction of merged puncta over total TMEM184B puncta area. **F,** Fraction of merged puncta over the total red pixel area of endosomal markers. **G,** Fraction of merged puncta (TMEM184B + LAMP1 + CTSD) over total TMEM184B pixel area. **H,** Fraction of merged puncta over the total red pixel area (LAMP1 + CTSD) of lysosomal markers. Error bars represent SEM.

To assess lysosomal localization, we used antibodies against LAMP1 and CTSD. Notably, LAMP1 is not exclusively localized to lysosomes but is also present in late endosomes (Cheng et al., 2018; Yap et al., 2022; Yap & Winckler, 2022). Therefore, the addition of CTSD is required for accurate evaluation of lysosomes. We first quantified the overlap between TMEM184B^+^ and LAMP1^+^CTSD^+^ puncta relative to the total number of TMEM184B^+^ puncta in HEK293T cells. Notably, TMEM184B^+^ puncta exhibited minimal colocalization with the lysosomal markers LAMP1 and CTSD (Fig. 2G). However, when considering the total lysosomal population, approximately 24% of lysosomes contained TMEM184B, indicating its presence on a subset of lysosomes (Fig. 2H). These findings suggest that TMEM184B is broadly distributed throughout the endosomal system, with a greater enrichment in early endosomal compartments.

### TMEM184B loss reduces endolysosomal acidification

The interactions between TMEM184B and the V-ATPase (Fig. 1D and 1-2) suggest that TMEM184B may regulate V-ATPase function. The endosomal system comprises a dynamic network of compartments that undergo progressive acidification during maturation (Hu et al., 2015; Wang et al., 2017). We focused on TMEM184B’s role in endolysosomal acidification because of recent attention towards lysosomal deacidification in neurodegenerative disorders and aging (Boland et al., 2008; Colacurcio & Nixon, 2016; J. H. Lee et al., 2022; Lie & Nixon, 2019; Nixon & Yang, 2012; Wolfe et al., 2013). To explore this, we utilized FIRE-pHly (Chin et al., 2021), a pH-sensitive sensor tagged to LAMP1, to assess the acidity of these compartments in primary embryonic cortical neurons from *Tmem184b*-mutant and wild-type mice. Changes in compartment acidification were assessed by the ratio of green fluorescence (mTFP1) to red fluorescence (mCherry) in FIRE-pHly^+^ puncta.

FIRE-pHly^+^ puncta are primarily localized near the soma and initial projection segments, as expected for lysosomes (Fig 3A-B). Notably, we observed that *Tmem184b*-mutant neurons occasionally displayed large puncta with prominent green fluorescence (Fig. 3B). We suspect these abnormal, enlarged compartments represent deacidified late endosomes or lysosomes, as wild-type neurons did not display similarly shaped puncta. In the absence of TMEM184B, FIRE-pHly^+^ puncta exhibited a significantly increased average green-to-red (G/R) fluorescence ratio in each neuron (p = 0.0189), indicating deacidification of these compartments compared to wild-type controls (Fig. 3C). *Tmem184b*-mutant neurons exhibit deacidified puncta regardless of size (Extended Data Figure 3-1). Average puncta size showed no significant difference between *Tmem184b-*mutant neurons and wild-type controls (Fig. 3D). Furthermore, no significant change in the density of FIRE-pHly^+^ puncta was observed (Fig. 3E). Together, this data shows that TMEM184B is necessary for proper acidification of the endolysosomal system.

**Fig 3:**
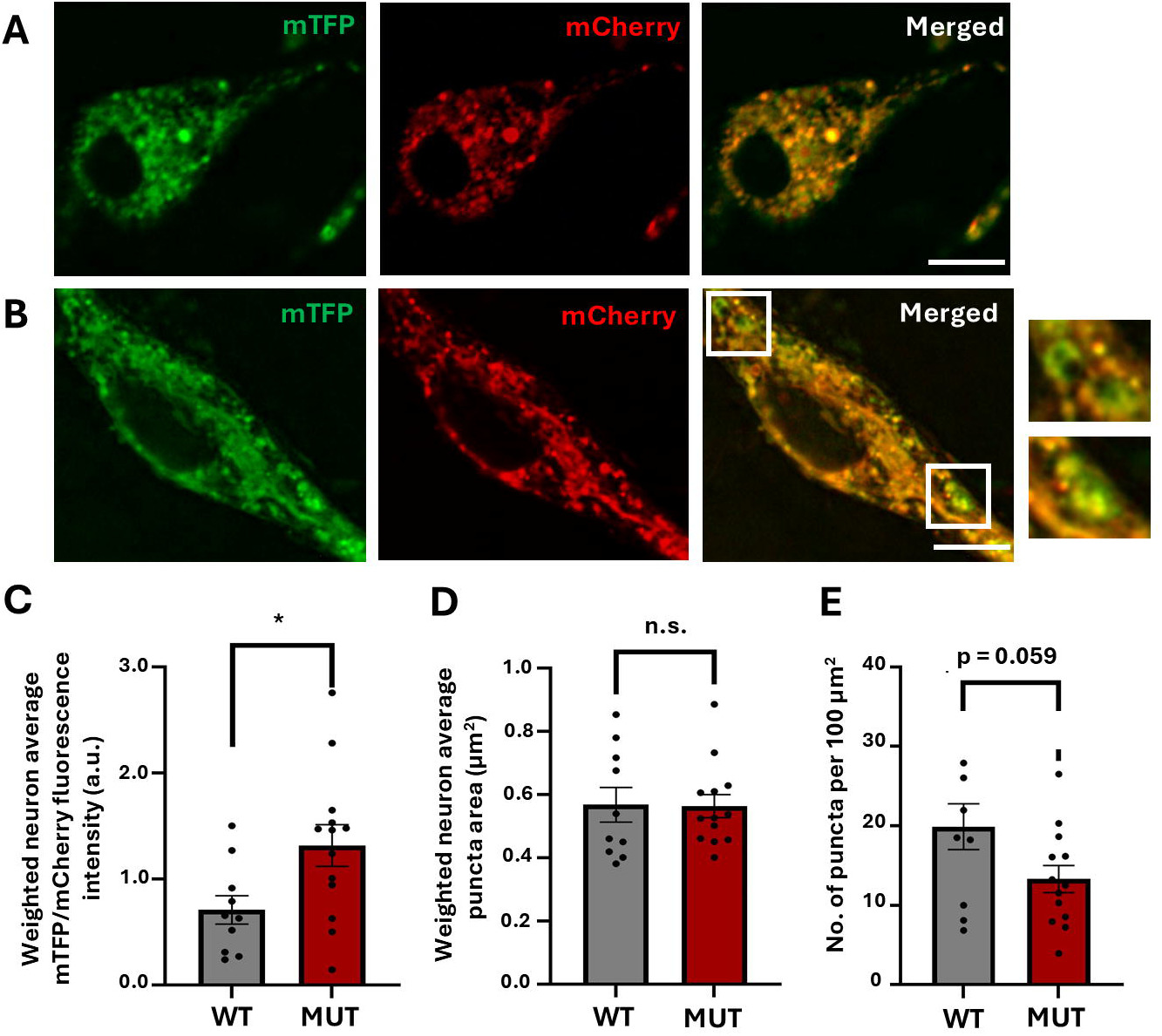
TMEM184B loss perturbs endolysosomal acidification. All images are single slices of the cell body and initial projections. **A-B,** Cell bodies and initial segments from wild-type (A) and *Tmem184b*-mutant (B) embryonic cortical neurons transfected with FIRE-pHly. Cropped images depict enlarged, abnormal green puncta indicated with white boxes. Scale bars = 5 µm. **C,** Comparison of weighted wild-type and mutant average G/R fluorescence intensity (p = 0.0189) using unpaired t-test with Welch’s correction. **D,** Weighted average puncta area (µm^2^) of wild-type and mutant neurons (p = 0.751) using Mann Whitney test. **E,** Number of puncta normalized to the cell body area per 100 μm^2^ for each neuron (p = 0.059). Using unpaired t-test with Welch’s correction. Error bars represent SEM.

### Loss of TMEM184B disrupts assembly of the vesicular proton pump

V-ATPase activity is regulated by the assembly of its V0 and V1 subcomplexes. The formation of these complexes is influenced by lumenal nutrient levels and growth signaling pathways, adjusting to the cell’s metabolic needs. As we saw reduced endolysosomal acidification in the absence of TMEM184B, we wanted to determine if this reduction was due to reduced V-ATPase assembly through co-IP assays. Our results revealed significantly reduced interactions between the subunit V0a1 and both V1A and V1H subunits in *Tmem184b*-mutant mice (Fig. 4A-B) relative to wild-type controls. The interaction between V0a1 and V1H was reduced by 31.72% in the absence of TMEM184B, while the interaction between V0a1 and V1A was reduced by 64% (Fig. 4C, E). Quantification of total V1H and V1A levels in lysates indicated that the reduced interaction was not due to a decrease in protein expression (Fig. 4D, F). These findings indicate that the absence of TMEM184B disrupts V-ATPase assembly, which offers a plausible mechanistic explanation for the decreased endolysosomal acidification seen in *Tmem184b*-mutant neurons.

**Fig 4:**
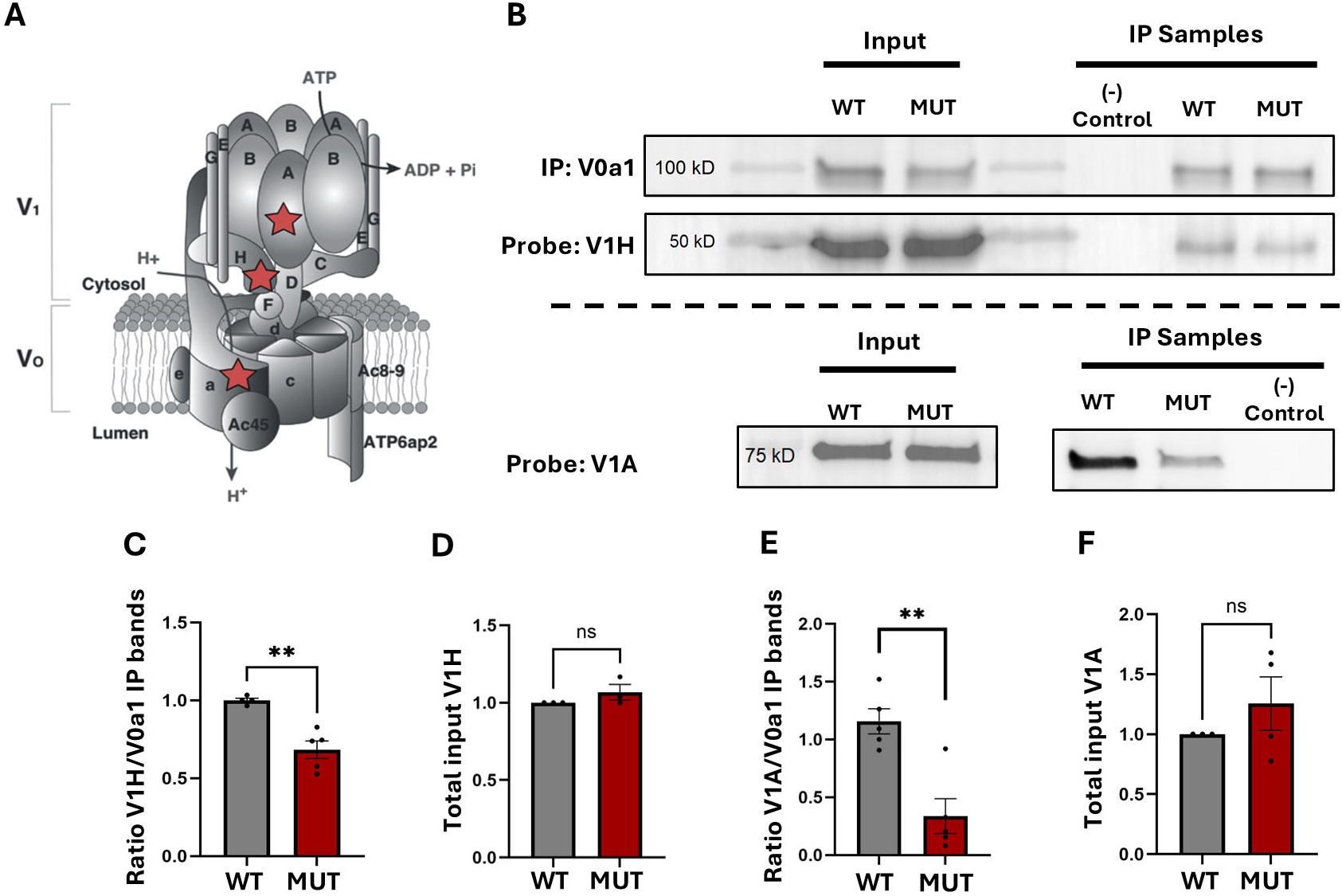
Loss of TMEM184B disrupts assembly of the vesicular proton pump. **A**, Diagram of the vesicular proton pump adapted from previous work (Sun-Wada & Wada, 2015). Red stars indicate specific subunits targeted in assembly assay. **B,** Blots of mouse hippocampal lysates show presence of V1H (top) and V1A (bottom) after V0a1 pulldown in wild-type and mutant mice. **C and E,** Quantification of the interaction between V0 and V1 subunits in *Tmem184b-*mutants and wild-type controls. Results of IP analysis normalized to the average total of V0A1 band intensity in wild-type IP samples. p = 0.0039 (C), p = 0.0028 (E). **D and F,** Quantification of total V1H (D) and V1A (F) normalized to the total amount of V0a1 input for IP. p = 0.32 (D), p = 0.33 (F). For all graphs, statistical evaluation used unpaired t-test with Welch’s correction. Error bars represent SEM.

## Discussion

Neurons are highly polarized cells with complex morphologies, characterized by multiple elongated projections extending from the cell soma. As post-mitotic cells, neurons encounter unique challenges in preserving their structure and function over their extended lifespan. Endolysosomal trafficking plays an essential role in supporting neuronal growth and long-term maintenance by facilitating the proper localization of cellular proteins to membranes, regulating intracellular signaling pathways, and coordinating the clearance of endocytosed cargo (Kuijpers et al., 2021; Roney et al., 2022). Dysregulated endolysosomal trafficking, particularly due to compartmental deacidification, contributes to cellular dysfunction associated with many neurological disorders. Despite significant progress, the mechanisms by which neurons maintain endolysosomal pH remain incompletely understood. Elucidating these mechanisms may reveal critical therapeutic targets for preventing the accumulation of toxic proteins and mitigating neurodegeneration.

Our data indicate that TMEM184B interacts with candidate proteins associated with endosomal trafficking and macroautophagy. Among these were the V0a subunit (a1 and a2 isoforms) of the V-ATPase. Evidence has shown that a1 resides on lysosomes and endosomes while a2 is present on the Golgi and early endosomes (Tuli & Kane, 2023). We also found an association with the RAVE complex and DMXL1 (Rabconnectin-3a), both well-established regulators of V-ATPase complex assembly. These findings prompted us to investigate the hypothesis that TMEM184B is not only localized within the endosomal system beyond that of recycling endosomes but also that TMEM184B plays a role in regulating endosomal acidification.

TMEM184B has not been identified before in association studies that defined the proteins contributing to the V-ATPase subcomplexes or their regulatory proteins (Merkulova et al., 2015). This could be because TMEM184B is not highly expressed (Bhattacharya et al., 2016; Larsen et al., 2022) and therefore may have fallen under the threshold for inclusion as a true positive interactor in directed immunoprecipitation or in high-throughput studies of proteome-wide interactions. In fact, the most highly cited association study acknowledges that their thresholds were set high and thus true interactors at low abundances may not be identified (Merkulova et al., 2015). Because we wanted to ensure our IP-MS results were robust, we used strict filtering parameters to identify positive interacting candidates. Although our interaction data is derived from over-expression models of TMEM184B, we observed a robust and bi-directional interaction between TMEM184B and the V0a1 subunit (Fig. 1-2).

In human cells, TMEM184B is localized to early endosomes, late endosomes and recycling endosomes (Fig. 2). During the transition from early to late endosomes, a stepwise process occurs in which Rab7 replaces Rab5 on the endosomal membrane (Langemeyer et al., 2020; Mottola, 2014; Skjeldal et al., 2021). Therefore, both Rab GTPases can transiently coexist on the same compartment (van der Beek et al., 2022). This may explain the high fractions of TMEM184B puncta overlapping with Rab5 or Rab7. In contrast, very few TMEM18B puncta are localized to lysosomes. These findings indicate that while TMEM184B localizes to both early and late endosomes, its primary function may be associated with the early stages of endocytosis rather than the terminal degradation phase. One possible explanation for this that TMEM184B is degraded in lysosomes or retained in late endosomes during late endosome-lysosome (LEL) fusion. If TMEM184B is sorted into intraluminal vesicles (ILVs) of late endosomes, it would be directed towards lysosomal degradation. Overexpression of TMEM184B may increase its degradation in lysosomes, consistent with the observation that approximately 20% of the total lysosomal population (LAMP1^+^CTSD^+^) colocalized with TMEM184B. TMEM184B exhibited partial localization at the plasma membrane, which may result from fusion with recycling endosomes rather than endogenous surface expression.

The absence of TMEM184B leads to reduced acidification of late endosomes and lysosomes in neurons (Fig. 3). As endosome maturation is tightly regulated by acidification, we hypothesized that these compartments’ size and morphology may also be perturbed. However, while we did occasionally observe enlarged and deacidified endolysosomes, we did not see a significant difference in the size of endolysosomes or the density of endolysosomes in the cell body. This suggests that the reduced endolysosomal acidification observed in mutant mice does not affect the morphology or distribution of degradative compartments within the cell body. Together, these results could suggest a perturbation in late endosome-lysosome fusion, leading to alkaline lysosomes residing in the cell body. An alternative explanation may be that impaired axonal retrograde late endosome transport reduces late endosome-lysosome fusion in the absence of TMEM184B. An interesting aspect to our findings is that TMEM184B appears to regulate lysosomal acidification without prominent lysosomal localization. It may achieve this by facilitating the budding and trafficking of the V0 subcomplex from the Golgi or endosomes, an essential step for V-ATPase assembly on endosomes and lysosomes. We identified interactors that participate in Golgi transport and vesicular tethering, suggesting that TMEM184B could participate early in the packaging or delivery of V-ATPase subunits from the Golgi throughout the endosomal system. Future studies should test these possibilities, which could illuminate how TMEM184B affects budding, fission, and fusion processes of membrane-bound compartments in neurons.

In this study, we demonstrate a reduction in V-ATPase subdomain assembly in the absence of TMEM184B. IP-MS analysis showed enriched associations between TMEM184B and two isoforms of the V0a subunit, a1 and a2 (Fig. 1). Given the observed reduction of endolysosomal acidification in TMEM184B-deficient cells (Fig. 3), we hypothesized that V-ATPase assembly might also be compromised. Supporting this hypothesis, we observed an approximately two-fold decrease in the interaction between the V0a1 and V1A subunits compared to V1H (Fig. 4), which was not attributable to reduced expression of either subunit. The V1H subunit is known to associate with the V0a1 subunit during complex assembly, and this substantial difference in interaction levels may reflect the spatial organization of these subunits. In the disassembled state, the V1H subunit undergoes a conformational change that brings it into closer contact with subunit F of the central stalk, effectively preventing ATP hydrolysis and rotational activity (Collins & Forgac, 2020). Future biochemical and structure-function studies may enable us to understand how TMEM184B influences pump function.

This study identifies a novel role for TMEM184B as a regulator of endolysosomal acidification. However, several aspects of its function remain unclear, including its precise biological role. The classification of TMEM184B within the transporter-opsin-G protein-coupled receptor (TOG) superfamily suggests a potential function as a transporter (Yee et al., 2013). We suspect that the functional role of TMEM184B may affect both localization to the appropriate intracellular compartments and proton pump activity of the V-ATPase, perhaps through separate and possibly indirect mechanisms. While de-orphanizing putative transporters is technically challenging, identifying the molecular substrates that may be transported by TMEM184B is of significant interest. Some possibilities consistent with our results are that TMEM184B could participate as an endosomal nutrient sensor for the V-ATPase or that it could indirectly modulate V-ATPase activity via ion counterbalance within endosomes. Such investigations are expected to refine the working model of TMEM184B’s functional role and would build upon our results to better understand and treat TMEM184B-associated neurodevelopmental disorders and other nervous system disorders featuring endolysosomal dysfunction.

## Data and Code Availability

All custom code developed for analysis will be freely available under an MIT license on Github at: https://github.com/eriklarsen4/

## Acknowledgements

The authors would like to acknowledge the support of the Nikon Imaging Center of Excellence at the University of Arizona. We thank Patty Jansma, UA Microscopy Core Facility, for early assistance with confocal microscopy and the staff of the Proteomics Shared Resource at the University of Arizona for assistance preparing sample for analysis. We thank Ross Buchan, Anita Koshy, Jean Wilson, and Konrad Zinsmaier and all members of the Bhattacharya lab for helpful discussions in the preparation of this manuscript. This work was funded by an NIH R01 (NS105680) to M.R.C.B.

---

**Extended Data Table 1-1:** Raw mass spectrometry data from IP-MS of TMEM184B (myc and V5-tagged) showing individual trial results, average values and fold changes of all proteins compared to controls.

**Extended Data Figure 1-2:** IP-MS validation of V-ATPase interaction with TMEM184B. FLAG blots split to optimize visualization of TMEM184B input bands compared to GFP input. **A,** Representative blots showing V0a1 content following FLAG pulldown in FLAG-TMEM184B and FLAG-GFP expressing cells. (n = 3 per group). **B**, Quantification of V0a1 bands normalized to the total level of FLAG-tagged protein in corresponding lane. **C,** Representative blots of FLAG-tagged protein content following V0a1 subunit pulldown in FLAG-TMEM184B and FLAG-GFP expressing cells. (n = 3 per group). **D**, Quantification of FLAG signal normalized to the total amount of V0a1 in corresponding lane. Error bars represent SEM.

**Extended Data Table 1-3:** Gene Ontology analysis results for the 136 proteins showing fold enrichment, adjusted p-values and false discovery fate (FDR) of biological process, cellular component and molecular function categories.

**Extended Data Figure 3-1:** Puncta acidity is consistently higher in *Tmem184b-*neurons regardless of puncta size. Yellow dots represent wild-type neurons. Blue dots represent *Tmem184b*-mutant neurons. **A**, Comparing distribution of individual puncta area (µm^2^) and corresponding G/R fluorescence intensity. **B,** Comparison of weighted average puncta area per neuron imaged with average G/R fluorescence ratio.

## References

Albishi, N. M. (2023). Autophagy genes AMBRA1 and ATG8 play key roles in midgut remodeling of the yellow fever mosquito, Aedes aegypti. Frontiers in Insect Science, 3. 10.3389/finsc.2023.1113871

Bar, M., Leibman, M., Schuster, S., Pitzhadza, H., & Avni, A. (2013). EHD1 Functions in Endosomal Recycling and Confers Salt Tolerance. PLOS ONE, 8(1), e54533. 10.1371/JOURNAL.PONE.0054533

Bhattacharya, M. R. C., Geisler, S., Pittman, S. K., Doan, R. A., Weihl, C. C., Milbrandt, J., & DiAntonio, A. (2016a). TMEM184b promotes axon degeneration and neuromuscular junction maintenance. Journal of Neuroscience, 36(17), 4681–4689. 10.1523/JNEUROSCI.2893-15.2016

Bhattacharya, M. R. C., Geisler, S., Pittman, S. K., Doan, R. A., Weihl, C. C., Milbrandt, J., & DiAntonio, A. (2016b). TMEM184b promotes axon degeneration and neuromuscular junction maintenance. Journal of Neuroscience, 36(17), 4681–4689. 10.1523/JNEUROSCI.2893-15.2016

Boland, B., Kumar, A., Lee, S., Platt, F. M., Wegiel, J., Yu, W. H., & Nixon, R. A. (2008). Autophagy induction and autophagosome clearance in neurons: Relationship to autophagic pathology in Alzheimer’s disease. Journal of Neuroscience, 28(27), 6926–6937. 10.1523/JNEUROSCI.0800-08.2008

Cataldo, A. M., Hamilton, D. J., Barnett, J. L., Paskevich, P. A., & Nixonls, R. A. (1996). Properties of the Endosomal-Lysosomal System in the Human Central Nervous System: Disturbances Maik Most Neurons in Populations at Risk to Degenerate in Alzheimer’s Disease. In The Journal of Neuroscience (Vol. 16, Issue 1).

Cataldo, A. M., Peterhoff, C. M., Troncoso, J. C., Gomez-Isla, T., Hyman, B. T., & Nixon, R. A. (2000). Endocytic Pathway Abnormalities Precede Amyloid Deposition in Sporadic Alzheimer’s Disease and Down Syndrome Differential Effects of APOE Genotype and Presenilin Mutations. In American Journal of Pathology (Vol. 157, Issue 1).

Chadwick, S. R., Grinstein, S., & Freeman, S. A. (2021). From the inside out: Ion fluxes at the centre of endocytic traffic. In Current Opinion in Cell Biology (Vol. 71, pp. 77–86). Elsevier Ltd. 10.1016/j.ceb.2021.02.006

Chapman, K., Yahiku, Z.A., Ullah F., Kodiparthi, S.V., Kellaris, G., Corriea, S.P., Stodberg, T., Sofokleous, C., Marinakis, N.M., Fryssira, H., Tsoutsou, E., Traeger-Synodinos, J., Accogli, A., Salpiero, V., Striano, S., Berger, S.I., Pond, K.W., Sirimulla, S., Davis, E.E., Bhattacharya, M.R.C. Pathogenic variants in TMEM184B cause a neurodevelopmental syndrome via alteration of metabolic signaling. medRxiv, 10.1101/2024.06.27.24309417v1.

Cheng, X. T., Xie, Y. X., Zhou, B., Huang, N., Farfel-Becker, T., & Sheng, Z. H. (2018). Characterization of LAMP1-labeled nondegradative lysosomal and endocytic compartments in neurons. Journal of Cell Biology, 217(9), 3127–3139. 10.1083/JCB.201711083

Chin, M. Y., Patwardhan, A. R., Ang, K. H., Wang, A. L., Alquezar, C., Welch, M., Nguyen, P. T., Grabe, M., Molofsky, A. V, Arkin, M. R., & Kao, A. W. (2021). Genetically Encoded, pH-Sensitive mTFP1 Biosensor for Probing Lysosomal pH. ACS Sensors, 6(6), 2168–2180. 10.1021/ACSSENSORS.0C02318

Cho, T. S., Beigaitė, E., Klein, N. E., Sweeney, S. T., & Bhattacharya, M. R. C. (2022). The Putative Drosophila TMEM184B Ortholog Tmep Ensures Proper Locomotion by Restraining Ectopic Firing at the Neuromuscular Junction. Molecular Neurobiology. 10.1007/S12035-022-02760-3

Choi, H., Larsen, B., Lin, Z. Y., Breitkreutz, A., Mellacheruvu, D., Fermin, D., Qin, Z. S., Tyers, M., Gingras, A. C., & Nesvizhskii, A. I. (2010). SAINT: probabilistic scoring of affinity purification–mass spectrometry data. Nature Methods 2010 8:1, 8(1), 70–73. 10.1038/nmeth.1541

Choudhury, A., Dominguez, M., Puri, V., Sharma, D. K., Narita, K., Wheatley, C. L., Marks, D. L., & Pagano, R. E. (2002). Rab proteins mediate Golgi transport of caveola-internalized glycosphingolipids and correct lipid trafficking in Niemann-Pick C cells. Journal of Clinical Investigation, 109(12), 1541–1550. 10.1172/jci200215420

Colacurcio, D. J., & Nixon, R. A. (2016). Disorders of lysosomal acidification—The emerging role of v-ATPase in aging and neurodegenerative disease. In Ageing Research Reviews (Vol. 32, pp. 75–88). Elsevier Ireland Ltd. 10.1016/j.arr.2016.05.004

Collins, M. P., & Forgac, M. (2020). Regulation and function of V-ATPases in physiology and disease. In Biochimica et Biophysica Acta - Biomembranes (Vol. 1862, Issue 12). Elsevier B.V. 10.1016/j.bbamem.2020.183341

Cotter, K., Stransky, L., McGuire, C., & Forgac, M. (2015). Recent Insights into the Structure, Regulation, and Function of the V-ATPases. In Trends in Biochemical Sciences (Vol. 40, Issue 10, pp. 611–622). Elsevier Ltd. 10.1016/j.tibs.2015.08.005

Di Bartolomeo, S., Corazzari, M., Nazio, F., Oliverio, S., Lisi, G., Antonioli, M., Pagliarini, V., Matteoni, S., Fuoco, C., Giunta, L., D’Amelio, M., Nardacci, R., Romagnoli, A., Piacentini, M., Cecconi, F., & Fimia, G. M. (2010). The dynamic interaction of AMBRA1 with the dynein motor complex regulates mammalian autophagy. Journal of Cell Biology, 191(1), 155–168. 10.1083/jcb.201002100

Eaton, A. F., Danielson, E. C., Capen, D., Merkulova, M., & Brown, D. (2024). Dmxl1 Is an Essential Mammalian Gene that Is Required for V-ATPase Assembly and Function In Vivo. Function, 5(4). 10.1093/function/zqae025

Forgac, M. (2007). Vacuolar ATPases: Rotary proton pumps in physiology and pathophysiology. In Nature Reviews Molecular Cell Biology (Vol. 8, Issue 11, pp. 917–929). 10.1038/nrm2272

Hu, Y. B., Dammer, E. B., Ren, R. J., & Wang, G. (2015). The endosomal-lysosomal system: From acidification and cargo sorting to neurodegeneration. In Translational Neurodegeneration (Vol. 4, Issue 1). BioMed Central Ltd. 10.1186/s40035-015-0041-1

Hurtado-Lorenzo, A., Skinner, M., Annan, J. El, Futai, M., Sun-Wada, G.-H., Bourgoin, S., Casanova, J., Wildeman, A., Bechoua, S., Ausiello, D. A., Brown, D., & Marshansky, V. (2006). V-ATPase interacts with ARNO and Arf6 in early endosomes and regulates the protein degradative pathway. Nature Cell Biology, 8(2), 124–136. 10.1038/ncb1348

Jaskolka, M. C., Tarsio, M., Smardon, A. M., Khan, M. M., & Kane, P. M. (2021). Defining steps in RAVE-catalyzed V-ATPase assembly using purified RAVE and V-ATPase subcomplexes. Journal of Biological Chemistry, 296. 10.1016/J.JBC.2021.100703

Jaskolka, M. C., Winkley, S. R., & Kane, P. M. (2021). RAVE and Rabconnectin-3 Complexes as Signal Dependent Regulators of Organelle Acidification. In Frontiers in Cell and Developmental Biology (Vol. 9). Frontiers Media S.A. 10.3389/fcell.2021.698190

Kane, P. M. (n.d.). Targeting Reversible Disassembly as a Mechanism of Controlling V-ATPase Activity.

Kruse, R., Krantz, J., Barker, N., Coletta, R. L., Rafikov, R., Luo, M., Højlund, K., Mandarino, L. J., & Langlais, P. R. (2017). Characterization of the CLASP2 Protein Interaction Network Identifies SOGA1 as a Microtubule-Associated Protein. Molecular & Cellular Proteomics : MCP, 16(10), 1718. 10.1074/MCP.RA117.000011

Kuijpers, M., Azarnia Tehran, D., Haucke, V., & Soykan, T. (2021). The axonal endolysosomal and autophagic systems. In Journal of Neurochemistry (Vol. 158, Issue 3, pp. 589–602). John Wiley and Sons Inc. 10.1111/jnc.15287

Langemeyer, L., Borchers, A. C., Herrmann, E., Füllbrunn, N., Han, Y., Perz, A., Auffarth, K., Kümmel, D., & Ungermann, C. (2020). A conserved and regulated mechanism drives endosomal rab transition. ELife, 9. 10.7554/eLife.56090

Larsen, E. G., Cho, T. S., McBride, M. L., Feng, J., Manivannan, B., Madura, C., Klein, N. E., Wright, E. B., Wickstead, E. S., Garcia-Verdugo, H. D., Jarvis, C., Khanna, R., Hu, H., Largent-Milnes, T. M., & Bhattacharya, M. R. C. (2022). Transmembrane protein TMEM184B is necessary for interleukin-31-induced itch. Pain, 163(5), E642–E653. 10.1097/j.pain.0000000000002452

Lee, J. H., Yang, D. S., Goulbourne, C. N., Im, E., Stavrides, P., Pensalfini, A., Chan, H., Bouchet-Marquis, C., Bleiwas, C., Berg, M. J., Huo, C., Peddy, J., Pawlik, M., Levy, E., Rao, M., Staufenbiel, M., & Nixon, R. A. (2022). Faulty autolysosome acidification in Alzheimer’s disease mouse models induces autophagic build-up of Aβ in neurons, yielding senile plaques. Nature Neuroscience, 25(6), 688–701. 10.1038/s41593-022-01084-8

Lee, J.-H., Yang, D.-S., Goulbourne, C. N., Im, E., Stavrides, P., Pensalfini, A., Chan, H., Bouchet-Marquis, C., Bleiwas, C., Berg, M. J., Huo, C., Peddy, J., Pawlik, M., Levy, E., Rao, M., Staufenbiel, M., & Nixon, R. A. (2022). Faulty autolysosome acidification in Alzheimer’s disease mouse models induces autophagic build-up of Aβ in neurons, yielding senile plaques. Nature Neuroscience, 25(6), 688–701. 10.1038/s41593-022-01084-8

Li, X., Lyu, Y., Li, J., & Wang, X. (2022). AMBRA1 and its role as a target for anticancer therapy. In Frontiers in Oncology (Vol. 12). Frontiers Media S.A. 10.3389/fonc.2022.946086

Lie, P. P. Y., & Nixon, R. A. (2019). Lysosome trafficking and signaling in health and neurodegenerative diseases. In Neurobiology of Disease (Vol. 122, pp. 94–105). Academic Press Inc. 10.1016/j.nbd.2018.05.015

Lloyd-Evans, E., Morgan, A. J., He, X., Smith, D. A., Elliot-Smith, E., Sillence, D. J., Churchill, G. C., Schuchman, E. H., Galione, A., & Platt, F. M. (2008). Niemann-Pick disease type C1 is a sphingosine storage disease that causes deregulation of lysosomal calcium. Nature Medicine, 14(11), 1247–1255. 10.1038/nm.1876

Lo, C. H., & Zeng, J. (2023). Defective lysosomal acidification: a new prognostic marker and therapeutic target for neurodegenerative diseases. Translational Neurodegeneration, 12(1), 29. 10.1186/s40035-023-00362-0

Mellacheruvu, D., Wright, Z., Couzens, A. L., Lambert, J. P., St-Denis, N. A., Li, T., Miteva, Y. V., Hauri, S., Sardiu, M. E., Low, T. Y., Halim, V. A., Bagshaw, R. D., Hubner, N. C., Al-Hakim, A., Bouchard, A., Faubert, D., Fermin, D., Dunham, W. H., Goudreault, M., … Nesvizhskii, A. I. (2013). The CRAPome: a contaminant repository for affinity purification–mass spectrometry data. Nature Methods 2013 10:8, 10(8), 730–736. 10.1038/nmeth.2557

Merkulova, M., Paunescu, T. G., Azroyan, A., Marshansky, V., Breton, S., & Brown, D. (2015). Mapping the H+ (V)-ATPase interactome: Identification of proteins involved in trafficking, folding, assembly and phosphorylation. Scientific Reports, 5. 10.1038/srep14827

Mottola, G. (2014). The complexity of Rab5 to Rab7 transition guarantees specificity of pathogen subversion mechanisms. Frontiers in Cellular and Infection Microbiology, 4(DEC). 10.3389/fcimb.2014.00180

Nardo, A. Di, Wertz, M. H., Kwiatkowski, E., Tsai, P. T., Leech, J. D., Greene-Colozzi, E., Goto, J., Dilsiz, P., Talos, D. M., Clish, C. B., Kwiatkowski, D. J., & Sahin, M. (2014). Neuronal Tsc1/2 complex controls autophagy through AMPK-dependent regulation of ULK1. Human Molecular Genetics, 23(14), 1–10. 10.1093/hmg/ddu101

Ng, S., Wu, Y. T., Chen, B., Zhou, J., & Shen, H. M. (2011). Impaired autophagy due to constitutive mTOR activation sensitizes TSC2-null cells to cell death under stress. Autophagy, 7(10), 1173–1186. 10.4161/auto.7.10.16681

Nixon, R. A. (2017). Amyloid precursor protein & endosomal-lysosomal dysfunction in Alzheimer’s disease: Inseparable partners in a multifactorial disease. FASEB Journal, 31(7), 2729–2743. 10.1096/fj.201700359

Nixon, R. A., & Rubinsztein, D. C. (2024). Mechanisms of autophagy–lysosome dysfunction in neurodegenerative diseases. Nature Reviews Molecular Cell Biology. 10.1038/s41580-024-00757-5

Nixon, R. A., & Yang, D. S. (2012). Autophagy and neuronal cell death in neurological disorders. Cold Spring Harbor Perspectives in Biology, 4(10). 10.1101/cshperspect.a008839

Parker, S. S., Krantz, J., Kwak, E. A., Barker, N. K., Deer, C. G., Lee, N. Y., Mouneimne, G., & Langlais, P. R. (2019). Insulin Induces Microtubule Stabilization and Regulates the Microtubule Plus-end Tracking Protein Network in Adipocytes. Molecular & Cellular Proteomics : MCP, 18(7), 1363. 10.1074/MCP.RA119.001450

Ratto, E., Chowdhury, S. R., Siefert, N. S., Schneider, M., Wittmann, M., Helm, D., & Palm, W. (2022). Direct control of lysosomal catabolic activity by mTORC1 through regulation of V-ATPase assembly. Nature Communications, 13(1). 10.1038/s41467-022-32515-6

Roney, J. C., Cheng, X. T., & Sheng, Z. H. (2022). Neuronal endolysosomal transport and lysosomal functionality in maintaining axonostasis. In Journal of Cell Biology (Vol. 221, Issue 3). Rockefeller University Press. 10.1083/jcb.202111077

Root, J., Merino, P., Nuckols, A., Johnson, M., & Kukar, T. (2021). Lysosome dysfunction as a cause of neurodegenerative diseases: Lessons from frontotemporal dementia and amyotrophic lateral sclerosis. In Neurobiology of Disease (Vol. 154). Academic Press Inc. 10.1016/j.nbd.2021.105360

Sardiu, M. E., Cai, Y., Jin, J., Swanson, S. K., Conaway, R. C., Conaway, J. W., Florens, L., & Washburn, M. P. (2008). Probabilistic assembly of human protein interaction networks from label-free quantitative proteomics. Proceedings of the National Academy of Sciences of the United States of America, 105(5), 1454. 10.1073/PNAS.0706983105

Singla, A., Fedoseienko, A., Giridharan, S. S. P., Overlee, B. L., Lopez, A., Jia, D., Song, J., Huff-Hardy, K., Weisman, L., Burstein, E., & Billadeau, D. D. (2019). Endosomal PI(3)P regulation by the COMMD/CCDC22/CCDC93 (CCC) complex controls membrane protein recycling. Nature Communications 2019 10:1, 10(1), 1–17. 10.1038/s41467-019-12221-6

Skjeldal, F. M., Haugen, L. H., Mateus, D., Frei, D. M., Rødseth, A. V., Hu, X., & Bakke, O. (2021). De novo formation of early endosomes during Rab5-to-Rab7a transition. Journal of Cell Science, 134(8). 10.1242/jcs.254185

Smardon, A. M., Tarsio, M., & Kane, P. M. (2002). The RAVE complex is essential for stable assembly of the yeast V-ATpase. Journal of Biological Chemistry, 277(16), 13831–13839. 10.1074/jbc.M200682200

Sowa, M. E., Bennett, E. J., Gygi, S. P., & Harper, J. W. (2009). Defining the Human Deubiquitinating Enzyme Interaction Landscape. Cell, 138(2), 389. 10.1016/J.CELL.2009.04.042

Sun-Wada, G. H., & Wada, Y. (2015). Role of vacuolar-type proton ATPase in signal transduction. Biochimica et Biophysica Acta (BBA) - Bioenergetics, 1847(10), 1166–1172. 10.1016/J.BBABIO.2015.06.010

Teo, G., Liu, G., Zhang, J., Nesvizhskii, A. I., Gingras, A. C., & Choi, H. (2013). SAINTexpress: improvements and additional features in Significance Analysis of Interactome software. Journal of Proteomics, 100, 37. 10.1016/J.JPROT.2013.10.023

Toei, M., Saum, R., & Forgac, M. (2010). Regulation and isoform function of the V-ATPases. In Biochemistry (Vol. 49, Issue 23, pp. 4715–4723). 10.1021/bi100397s

Tseng, C. C., Dean, S., Davies, B. A., Azmi, I. F., Pashkova, N., Payne, J. A., Staffenhagen, J., West, M., Piper, R. C., Odorizzi, G., & Katzmann, D. J. (2021). Bro1 stimulates vps4 to promote intralumenal vesicle formation during multivesicular body biogenesis. Journal of Cell Biology, 220(8). 10.1083/JCB.202102070/VIDEO-3

Tuli, F., & Kane, P. M. (2023). The cytosolic N-terminal domain of V-ATPase a-subunits is a regulatory hub targeted by multiple signals. In Frontiers in Molecular Biosciences (Vol. 10). Frontiers Media SA. 10.3389/fmolb.2023.1168680

van der Beek, J., de Heus, C., Liv, N., & Klumperman, J. (2022). Quantitative correlative microscopy reveals the ultrastructural distribution of endogenous endosomal proteins. Journal of Cell Biology, 221(1). 10.1083/jcb.202106044

Vivas, O., Tiscione, S. A., Dixon, R. E., Ory, D. S., & Dickson, E. J. (2019). Niemann-Pick Type C Disease Reveals a Link between Lysosomal Cholesterol and PtdIns(4,5)P2 That Regulates Neuronal Excitability. Cell Reports, 27(9), 2636–2648.e4. 10.1016/j.celrep.2019.04.099

Wang, C., Zhao, T., Li, Y., Huang, G., White, M. A., & Gao, J. (2017). Investigation of endosome and lysosome biology by ultra pH-sensitive nanoprobes. In Advanced Drug Delivery Reviews (Vol. 113, pp. 87–96). Elsevier B.V. 10.1016/j.addr.2016.08.014

Wolfe, D. M., Lee, J. hyun, Kumar, A., Lee, S., Orenstein, S. J., & Nixon, R. A. (2013). Autophagy failure in Alzheimer’s disease and the role of defective lysosomal acidification. European Journal of Neuroscience, 37(12), 1949–1961. 10.1111/ejn.12169

Wright, E. B., Larsen, E. G., Coloma-Roessle, C. M., Hart, H. R., & Bhattacharya, M. R. C. (2023). Transmembrane protein 184B (TMEM184B) promotes expression of synaptic gene networks in the mouse hippocampus. BMC Genomics, 24(1), 559. 10.1186/s12864-023-09676-9

Wyant, G. A., Abu-Remaileh, M., Wolfson, R. L., Chen, W. W., Freinkman, E., Danai, L. V, Vander Heiden, M. G., & Sabatini, D. M. (2017). mTORC1 Activator SLC38A9 Is Required to Efflux Essential Amino Acids from Lysosomes and Use Protein as a Nutrient. Cell, 171(3), 642–654.e12. 10.1016/j.cell.2017.09.046

Yap, C. C., Mason, A. J., & Winckler, B. (2022). Dynamics and distribution of endosomes and lysosomes in dendrites. In Current Opinion in Neurobiology (Vol. 74). Elsevier Ltd. 10.1016/j.conb.2022.102537

Yap, C. C., & Winckler, B. (2022). Spatial regulation of endosomes in growing dendrites. Developmental Biology, 486, 5–14. 10.1016/j.ydbio.2022.03.004

Yee, D. C., Shlykov, M. A., Västermark, Å., Reddy, V. S., Arora, S., Sun, E. I., & Saier, M. H. (2013). The transporter-opsin-G protein-coupled receptor (TOG) superfamily. FEBS Journal, 280(22), 5780–5800. 10.1111/febs.12499

Zhu, S., Quan, C., Wang, R., Liang, D., Su, S., Rong, P., Zhou, K., Yang, X., Chen, Q., Li, M., Du, Q., Zhang, J., Fang, L., Wang, H. Y., & Chen, S. (2022). The RalGAPα1–RalA signal module protects cardiac function through regulating calcium homeostasis. Nature Communications, 13(1), 4278. 10.1038/S41467-022-31992-Z

Zoncu, R., Bar-Peled, L., Efeyan, A., Wang, S., Sancak, Y., & Sabatini, D. M. (2011). mTORC1 senses lysosomal amino acids through an inside-out mechanism that requires the vacuolar H+-ATPase. Science, 334(6056), 678–683. 10.1126/SCIENCE.1207056

