## Supplementary figures and images for "Neuronal endolysosomal acidification relies on interactions between transmembrane protein 184B (TMEM184B) and the vesicular proton pump"

### Extended Data Figure 1-2

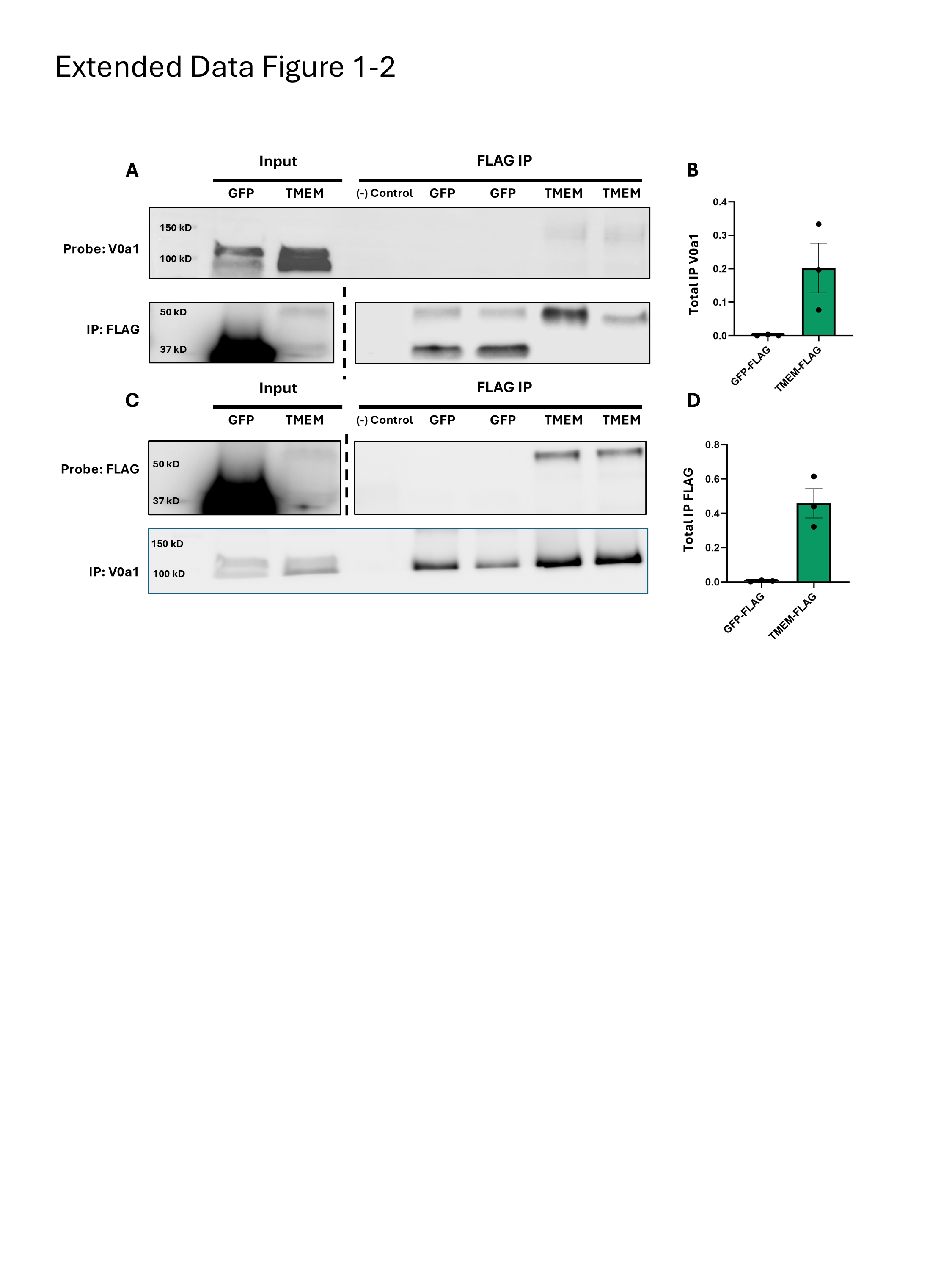

### Extended Data Figure 3-1

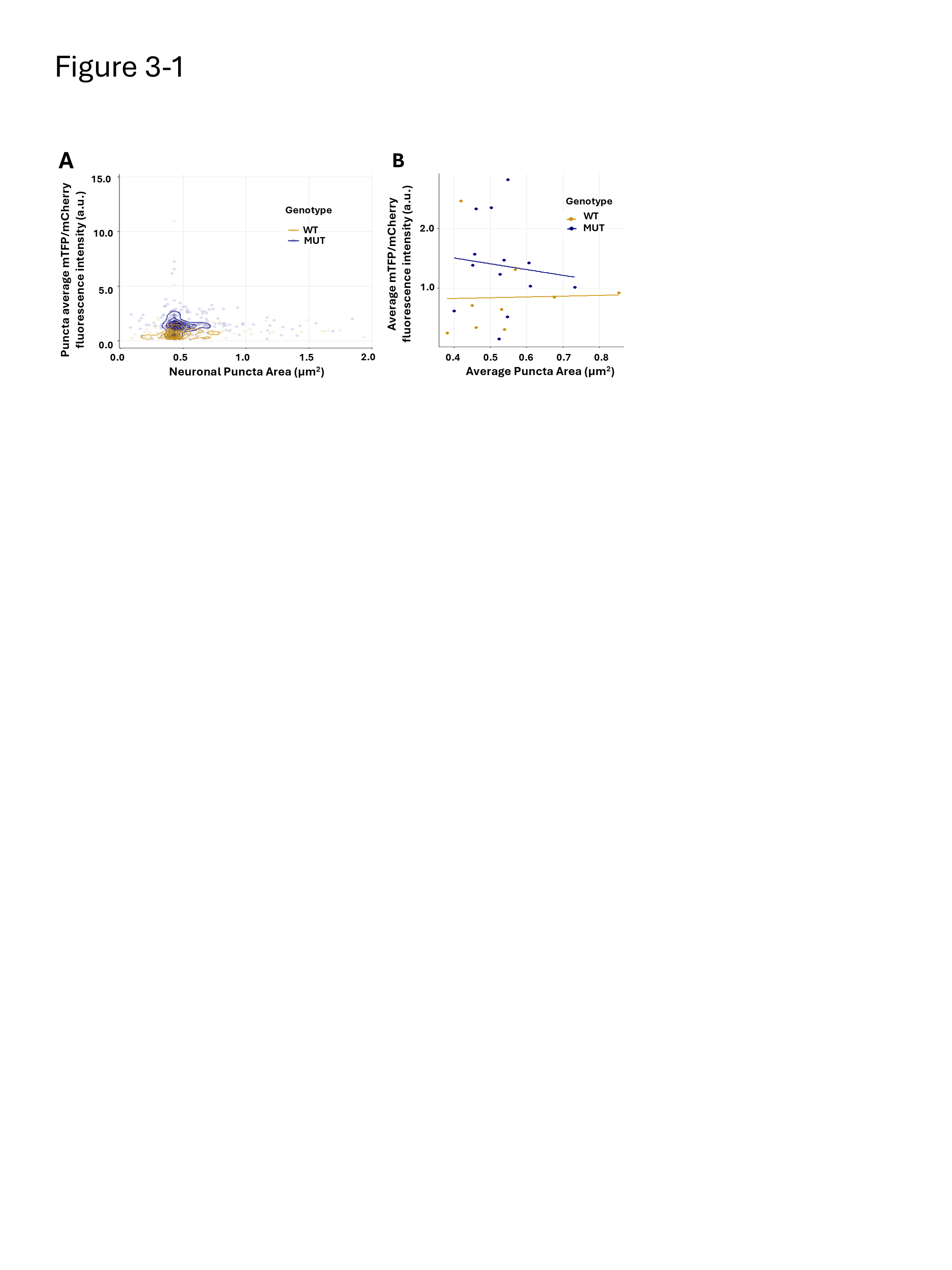
